# A Zur-mediated transcriptional regulation of the zinc export system

**DOI:** 10.1101/2022.09.30.510347

**Authors:** Verena Ducret, Diego Gonzalez, Sara Leoni, Martina Valentini, Karl Perron

## Abstract

The control of cellular zinc (Zn) concentrations by dedicated import and export systems is essential for the survival and virulence of *Pseudomonas aeruginosa*. The transcription of its many Zn transporters is therefore tightly regulated by a known set of transcription factors involved in either the import or the export of Zn. In this work, we show that the Zur protein, a well-known repressor of Zn import, plays a dual role and functions in both import and export processes. In a situation of Zn excess, Zur represses Zn entry, but also activates the transcription of *czcR*, a positive regulator of the Zn export system. To achieve this, Zur binds at two sites, located by DNA footprinting in the region downstream the *czcR* transcription start site. In agreement with this regulation, a delay in induction of the efflux system is observed in the absence of Zur and Zn resistance is affected. The discovery of this regulation highlights a new role of Zur as global regulator of Zn homeostasis in *P. aeruginosa* disclosing an important link between Zur and zinc export.

## Introduction

Metal ions are essential to the functioning and survival of all known forms of cellular life. Among all trace elements, zinc (Zn) is of particular importance because it functions as a cofactor of many essential enzymes and is, more generally, needed to sustain protein structure. Zn is found at the top of the Irving-Williams series (Mg < Ca < Mn < Fe < Co < Ni < Cu > Zn) that ranks metal ions according to their ability to form complexes with proteins (1). This is why, when present in excess inside the cell, Zn tends to outcompete other metals and cause protein mismetallization, the major cause of its cellular toxicity (2). The intracellular concentration of Zn has therefore to be tightly and finely controlled by complex homeostasis mechanisms. In bacteria, the cytoplasmic Zn concentration is maintained within the range of 10^-3^ to 10^-4^ M, making it the second most abundant metal after iron (3).

In the Gram-negative bacterium *Pseudomonas aeruginosa*, several systems are involved in Zn homeostasis (4,5). Zn import into the periplasm is mediated by TonB-dependent importers such as ZnuD. A P-type ATPase and three ABC transporters help with the uptake of Zn from the periplasm into the cytoplasm. In case of metal excess, Zn is exported by a specific P-type ATPase CadA, a Resistance-Nodulation-Cell Division (RND) efflux pump (CzcCBA), and two cation diffusion facilitators (CDF) that play a minor role in Zn resistance (3,6). Control of Zn import and export is extremely important for some bacterial pathogens, especially for their interaction with their host, whether an animal (7) or a plant (8). *P. aeruginosa*, for example, is able to grow and thrive in Zn-poor environments such as the bloodstream, where trace metals are chelated to prevent microbial growth (3). Conversely, in a phagolysosome, high concentrations of Cu and Zn have antimicrobial properties and can eliminate the pathogen (9). Several regulators of Zn homeostasis have been identified in *P. aeruginosa*. The major regulator of Zn import is the Zinc Uptake Regulator, hereafter Zur, protein (formerly Np20, (10)), a Fur (Ferric Uptake Regulator)-like protein (11). In the presence of an excess of cytoplasmic Zn, Zur binds to the metal ion, dimerizes, and, in its dimeric form, binds to specific Zur boxes on the DNA to repress genes involved in Zn import (4,10). Zn export is controlled by two other regulators: CadR, a MerR-type regulator that activates the transcription of the P-type ATPase CadA (12), and CzcR, a response regulator (RR) of the two-component system CzcRS, which, when phosphorylated by CzcS, activates the transcription of the efflux pump CzcCBA that expels Zn from the cell (13). Until now, the regulatory networks controlling Zn import and Zn export have been considered essentially independent from one another and no crosstalk between them has been substantiated.

In this study, we discovered that the Zur protein is needed for the full expression of the Zn efflux system when *P. aeruginosa* switches from a low-Zn environment to an excess-Zn situation. We show that Zur accelerates *czcR* transcription, and thus activates Zn efflux, by binding to two DNA sites in a region upstream the *czcR* translation start site (ATG). This new function makes Zur the keystone of Zn homeostasis in *P. aeruginosa*, acting not only as a repressor of Zn import but also as activator of Zn export.

## Results

### Zur binds to the czcRS promoter

The Zur protein, when concentrations of Zn are sufficient or excessive, binds to a 17 bp DNA Zur box to repress genes involved in Zn import (14). In a global bioinformatics search on the *P. aeruginosa* genome, we found, as expected, Zur boxes in the promoter regions of genes involved in the import of Zn, but also, intriguingly, in *czcRS* promoter region (5). This suggested a possible direct regulatory link, mediated by Zur, between the import and export of Zn. In order to validate the *in silico* analysis, we purified the Zur protein and performed an electrophoretic mobility shift assay (EMSA) on the *czcR* promoter. A clear DNA shift was observed in the presence of Zur (Fig. 1A). The interaction is Zn-dependent as no shift was detected in presence of TPEN, a Zn chelator. DNA footprint analysis revealed two boxes located approximately 40 bps upstream of the translation start codon of *czcR* (Fig 1B, 1C and S1A).

**Figure 1:**
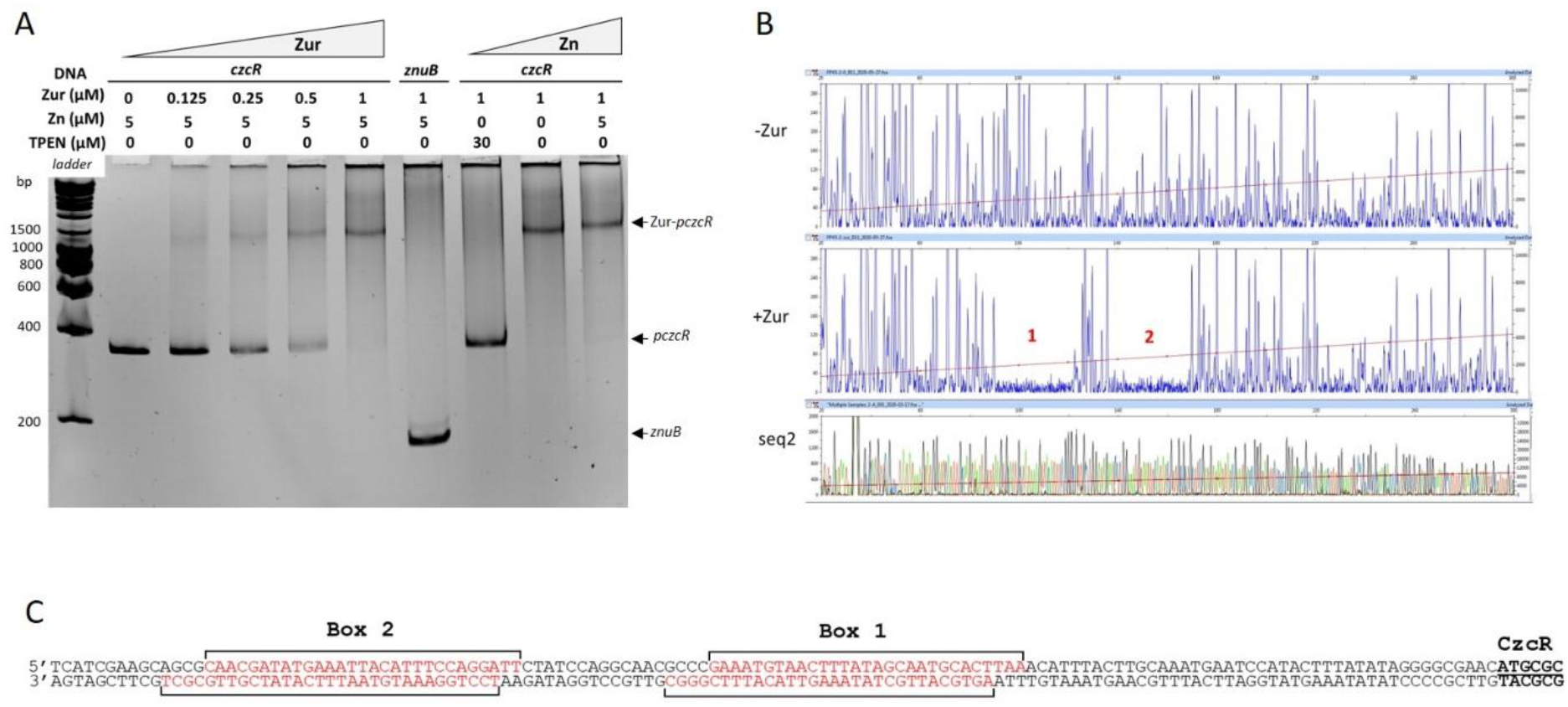
Zur binds upstream the *czcR* translation start codon. **A)**Electrophoretic Mobility Shift Assay (EMSA) using purified Zur protein and *czcR* or *znuB* DNA region as negative control. 25 ng of DNA per reaction was used. The Zur protein, Zn and TPEN (Zn chelator) concentrations are indicated at the top of the figure. **B)** DNAse I footprinting of *czcR* promoter in the absence or presence or of 1 μM Zur protein. DNA fragments were analyzed by capillary electrophoresis. The two Zur boxes are indicated by numbers 1 and 2 in red. The sequencing reaction is visible below the figure (seq2). C) Sequence of the two boxes as determined by the footprinting experiments (Fig 1B coding strand; Fig S1A template strand).

The transcription start site of *czcR* was determined by 5’RACE and it was found 300 bps upstream of the *czcR* translation start, revealing a 292-nucleotide 5’-UTR (Fig 2A, S1B). Promoter fusions (*-gfp*) confirmed that a region longer than 292 bps upstream of the ATG is required to sustain *czcRS* transcription in the presence of Zn (construct #1 and #2 in Fig 2B).

**Figure 2:**
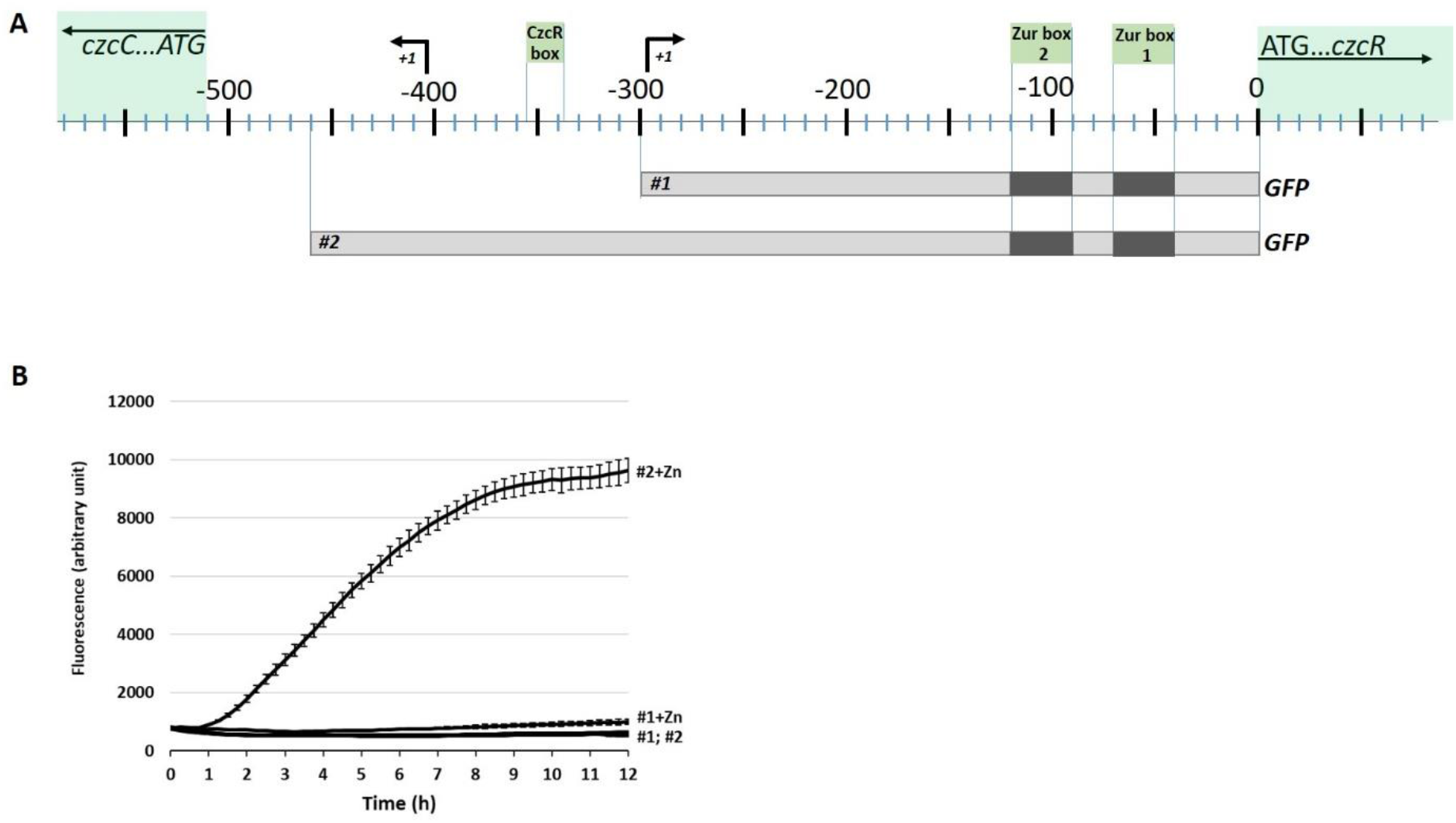
GFP reporter of the *czcR* promoters. **A)**Map of the *czcR* promoter and 5’-UTR region with the two Zur boxes, the CzcR DNA binding site (according to (15)) and the *czcR* and *czcC* transcription start sites (+1). The number represents the nucleotide position relative to the translation start of *czcR* gene. **B)**GFP promoter fusions #1 and #2 transformed into the wt PAO1 strain. Time 0 corresponds to the addition of 2 mM ZnCl2. Standard deviations of the triplicates are indicated.

To determine whether the binding at the two Zur boxes was independent, we repeated the band shift assay using different versions of the *czcR* promoter, where box1 (mut1), box 2 (mut2), or both (mut1+2) Zur boxes were mutated at key positions (Fig 3A). A higher shift was obtained when the two Zur boxes were present compared to the presence of only one, and no shift was detected when the two boxes were mutated (Fig 3B), indicating that the two boxes support the binding of the regulator independently.

**Figure 3:**
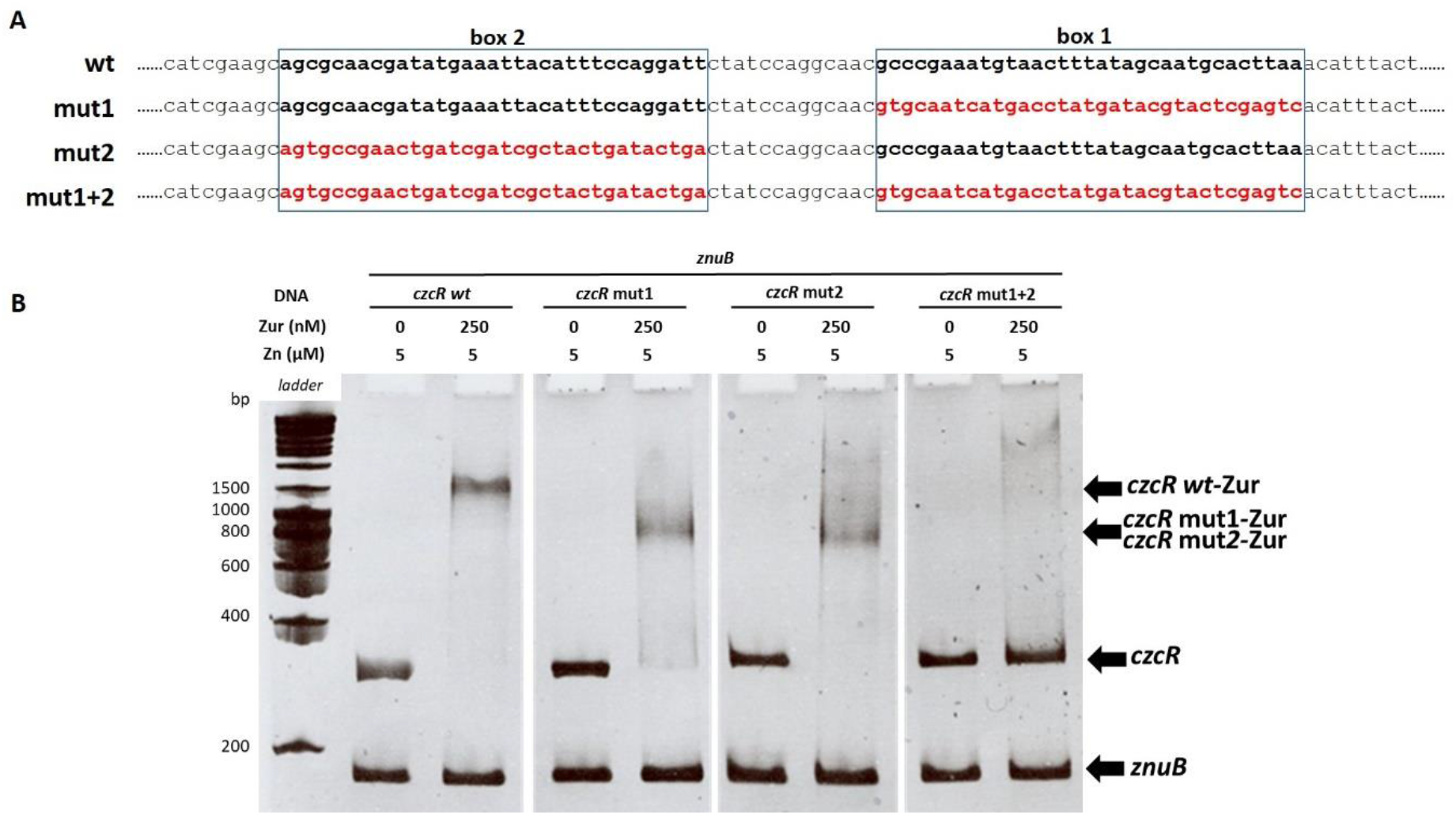
Mutations in the Zur boxes and their effects on Zur binding activity. **A)**Sequences of modified Zur boxes: mut1, mut2 and mut1+2 in red. Bold black boxes correspond to the wt sequence.**B)**Electrophoretic Mobility Shift Assay (EMSA) using purified Zur protein, different *czcR* 5’-UTR (*czcR*) and *znuB (*negative DNA control). 25 ng of each DNA per reaction was used. The Zur protein and Zn concentrations are indicated at the top of the figure. *czcR* wt, corresponds to the wt sequence of the *czcR* 5’-UTR, *czcR* mut1 and mut2 possess mutated box 1 or 2, respectively; *czcR* mut1+2 possess the two mutated Zur boxes.

### Zur is involved in zinc resistance

In order to investigate whether Zur had a role in the regulation of Zn export, we characterized the phenotype of a Δ*zur* mutant. We found that, in the absence of Zur, *P. aeruginosa* is more sensitive to Zn excess than the parental strain (Fig. 4A). The phenotype is milder than in a *ΔczcRS* mutant, but more severe than in a Δ*cadR* mutant (12).

**Figure 4:**
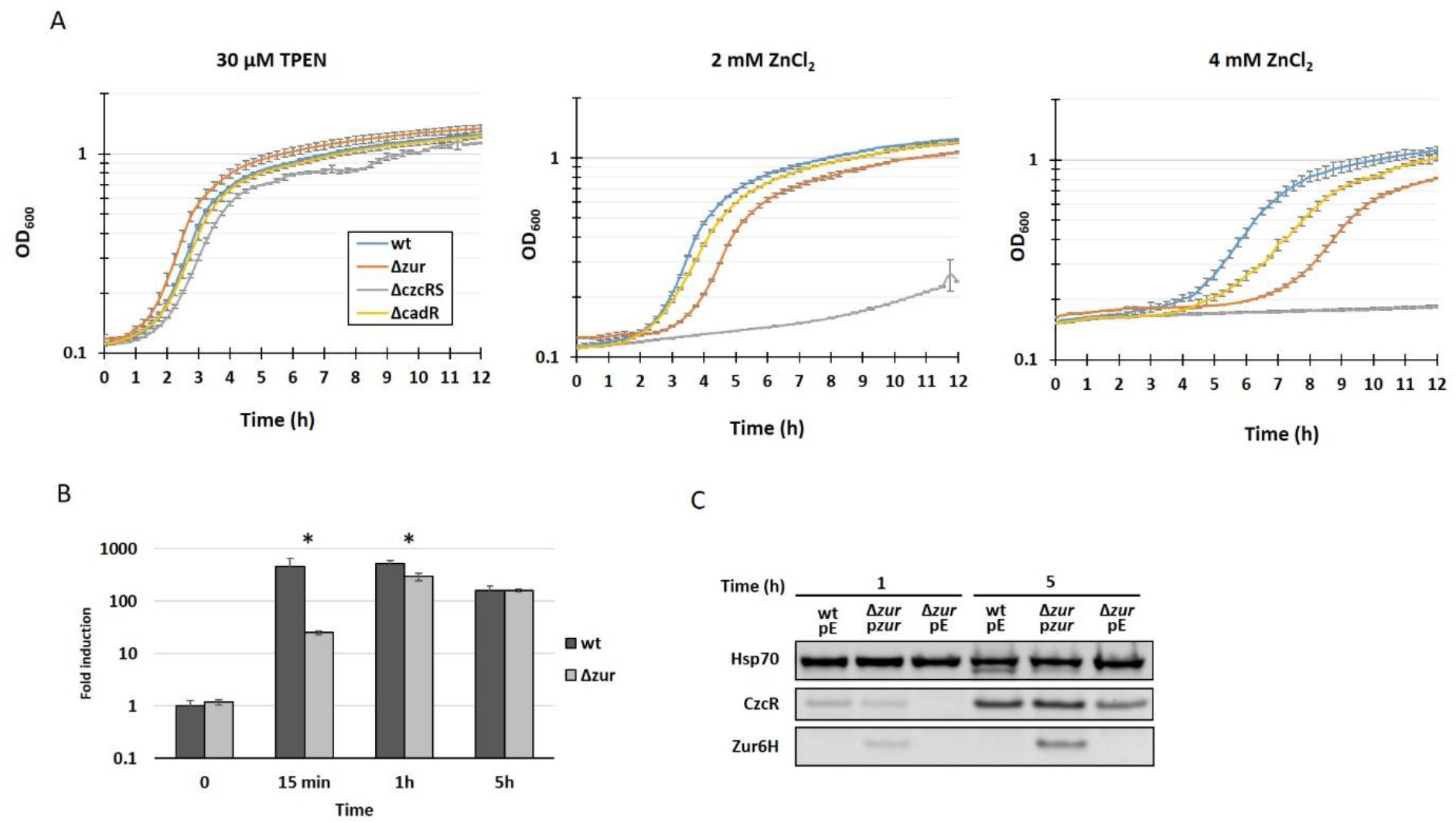
Zur is involved in CzcR expression. **A)**Growth curves of *P. aeruginosa* wt, *Δzur, ΔczcRS* and *ΔcadR* mutants in Zn-depleted medium (30 μM TPEN), in 2 mM and 4 mM ZnCl_2_, as indicated. **B)**Expression of *czcR* measured using qRT-PCR. RNAs were extracted at 0, 15 min, 1h and 5h after addition of 2 mM ZnCl_2_ as indicated. Fold expressions are compared to the wt strain at t0 and normalized by *oprF* expression. Standard deviations of the triplicates are indicated. Statistical analyses were performed according to the Student’s *t*-test and *p*-values are given as follows: ≤0.05 (*). **C)**Western blot analysis of total proteins sampled 1h and 5h after addition of 2 mM Zn. wt and Δ*zur* mutant containing the empty plasmid (pE) or the IPTG inducible *zur:6His* gene (p*zur*). Blots were decorated with anti-CzcR, anti-6His for Zur detection and anti-Hsp70 as loading control.

The effect of the *zur* deletion on *czcR* transcription was assessed using qRT-PCR. In the absence of Zur, *czcR* expression decreased significantly compared to the control (Fig 4B). The difference was obvious 15 minutes (400-fold decrease) after Zn addition, still visible after 1h (230-fold decrease), but no longer present after 5h of induction. An immunoblot analysis confirmed this expression pattern at the protein level (Fig 4C). As expected, the delay in *czcR* expression was abolished when *zur:6His* was overexpressed *in trans* by the addition of 0.2 mM IPTG (Fig 4C). In agreement with our model, the transcription of *czcCBA* and *czcD*, which are dependent on CzcR (4), was also affected in the *Δzur* mutant (Fig 5). Conversely, the P-type ATPase CadA, which is known to be overexpressed in the absence of CzcR (12), showed increased expression in the *Δzur* mutant (Fig 5). Altogether, these results strongly suggest that Zur is required to bring about the full expression of *czcR* and mount the proper response to a Zn excess. Deleting the Zur regulated major Zn import permease ZnuB has no effect on the *zur* mutant growth phenotype in the presence of Zn (Fig S2) suggesting a direct effect of Zur on *czcR* expression.

**Figure 5:**
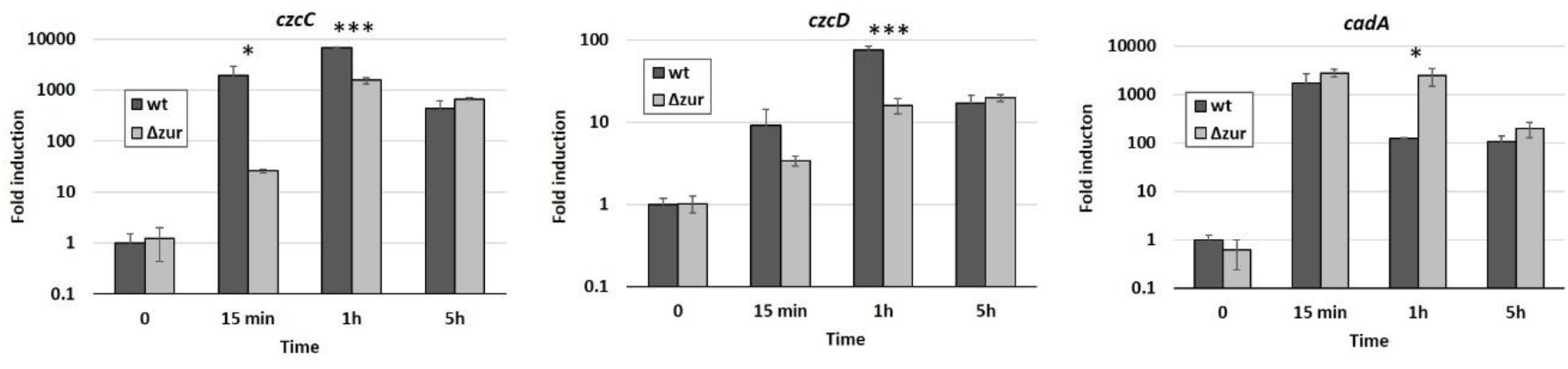
Expression of *czcC, czcD* and *cadA* measured using qRT-PCR. RNAs were extracted at 0, 15min, 1h and 5h after addition of 2 mM ZnCl_2_ as indicated. Fold expressions are compared to t0 and normalized by *oprF* expression. Standard deviations of the triplicates are indicated. Statistical analyses were performed according to the Student’s *t*-test and *p*-values are given as follows: ≤0.05 (*), ≤0.001 (***).

## Discussion

Zinc is an essential element whose cellular concentration is tightly regulated (16). It is vital for cells and must always be present in sufficient amounts, but can become toxic as soon as it exceeds a threshold (17). In the bacterium *P. aeruginosa*, eight zinc import and four zinc export systems have been described (4). In case of zinc excess, the Zur protein represses the expression of many import systems, thus preventing too much Zn from entering the cell. In this work we described a novel key function of the Zur protein in *P. aeruginosa*, being involved not only in the repression of the import of Zn, but also in the early and fast induction of the CzcCBA Zn export system. Zur is therefore a keystone of Zn homeostasis in *P. aeruginosa*, integrating both the import and export of the ion.

In some other bacteria, including *Xanthomonas campestris, Streptomyces coelicolor* and *Caulobacter crescentus*, Zur also acts as an activator of zinc efflux genes (18–20). The localization of the Zur boxes on the promoter region of its target genes, along with the ability of the protein to multimerize at high zinc concentrations, appear to determine whether the regulator functions as a repressor or an activator (6). Other processes, not directly related to metal homeostasis, have also been found to be positively regulated by Zur such as the synthesis of secondary metabolites in *Streptomyces avermitilis* or the type VI secretion system in *Yersinia pseudotuberculosis* (21,22). In the case of *P. aeruginosa*, we found that the Zur protein binds in a Zn-dependent manner to two DNA regions (Zur box1 and 2) located downstream of the *czcR* promoter. This binding has a positive effect on *czcR* transcription and, in general, for the Zur-mediated Zn resistance.

What is the molecular action of Zur for the activation of *czcR* transcription? Recently numerous TCS RRs have been detected in the intergenic region between *czcR* and *czcC* (23). Zur could therefore be involved in the competition with the binding of other transcriptional repressor in this region. Fine tuning of the efflux system could be essential in environmental conditions not tested in the laboratory. For instance, rapid changes in Zn concentration are known to occur during infection, because of nutritional immunity and metal excess in the phagolysosome (24,25). The tight efflux pump control might also be important in the environment, where rapid changes in metal concentrations might also occur. Interestingly, among the *Pseudomonas* species analyzed for the presence and location of Zur boxes (i.e. *P. chlororaphis, P. fluorescens, P. putida, P. stuzeri* and *P. syringae*), only *P. aeruginosa* possesses boxes upstream of the *czcR* gene (5). This characteristic could be related to its ability to proliferate in presence of high metal concentrations. Thus, in *P. aeruginosa*, Zur appears to be the central cog in Zn homeostasis since it directly controls both the import and export of this metal. CzcR is also known to be capable of controlling the expression of the gene coding for the OprD porin, involved in imipenem resistance, or the *lasI* gene involved in quorum sensing (13,26). The importance of Zur in the control of *czcR* expression highlights a novel and important role of Zur in the interaction between Zn homeostasis, virulence factor expression and antibiotic resistance in *P. aeruginosa*. Deciphering this novel and important regulation occurring by Zur at the *czcR* gene vicinity is therefore of prime interest.

## Materials and Methods

### Bacterial Strains and Culture Media

The bacterial strains are listed in supplementary table S1. The modified Luria-Bertani medium (M-LB) used for this work was prepared as mentioned previously (12). When required, antibiotics were added to the medium at the following concentrations: 50 μg/mL tetracycline (Tc, Axxora), 200 μg/mL carbenicillin (Cb, phytotechlab), 50 μg/mL Gentamycin (Gm, AppliChem) for *P. aeruginosa* or 100 μg/mL ampicillin (Ap, AppliChem) and 15 μg/mL Tc or Gm for *E. coli*.

### Genetic Manipulations

The primers used for cloning are detailed in supplementary table S2. Restriction and ligation enzymes (Promega), or fragment insertion using the Gibson assembly Cloning kit (New England Biolabs) were employed according to the supplier’s instructions. Plasmids were transformed into *E. coli* DH5α by heat shock, verified by sequencing before being electroporated into the *P. aeruginosa* wild type or *Δzur* strains (27). For transcriptional *gfp*-fusion assays, the two *czcR* promoter regions (#1 and #2 in Fig 2A) were obtained by PCR amplification. Fragments were then digested with KpnI and BglII enzymes and ligated into the corresponding sites of the pBRR1-*gfp* vector. For Zur6HIS overexpression assays, the full *zur* gene was amplified by PCR with a reverse primer containing the 6xHIS tag. The fragment was digested with the EcoRI and BamHI enzymes and ligated into the pMMB66EH plasmid. The pME6001-*czcRS* vector was used as a template for the sequencing reactions of the 5’RACE experiments.

For this construction, the full *czcRS* operon and its promoter region were amplified by PCR and cloned into the pME6001 vector with the XhoI and HindIII restriction sites. . For the *Δzur* mutant, a DNA product, consisting of the two 500 bp amplicons flanking the *zur* gene fused together, was obtained by megaprimer amplification, digested with EcoRI and BamHI restriction enzymes and ligated into the pME3087 plasmid according to standard procedure (28). After verification, the resulting plasmids were transformed into the *P. aeruginosa* wt strain. Merodiploids were selected as described previously (29) and deletions were confirmed by sequencing.

### Growth experiments

To monitor the growth of the wt and the mutant strains, overnight cultures were diluted in M-LB medium to an OD600 of 0.05, supplemented with N,N,N’,N’-tetrakis(2-pyridylmethyl)-ethylenediamine (TPEN; Brunschwig) or ZnCl_2_ (Fluka) as indicated in figures 3A and 4. Cultures were incubated at 37°C with shaking in a microplate reader (BioTek Instruments) and absorbance at 600 nm was measured every 15 min, for 12h. For strains carrying the pBRR1 plasmid with *gfp* fusions #1 or #2, overnight cultures were diluted to an OD600 of 0.1 and cultured for 2h30 in the presence of 30 μM TPEN before adding ZnCl_2_. Absorbance at 600 nm and fluorescence at 528 nm were then monitored every 15 min. Arbitrary units indicted in the figure 2B correspond to the fluorescence values normalized with the cell density.

### Immunoblot analyses

Immunoblot analyses were performed as previously described (26). Briefly, overnight cultures were diluted to an OD 0.1 and incubated 2h30 in M-LB supplemented with 30 μM TPEN. Isopropyl-1-thio-D-galactopyranoside (IPTG, Sigma Aldrich) was added at a final concentration of 0.2 mM to cultures complemented with Zur6HIS on the pMMB66EH or the empty plasmid. 1 ml was collected and centrifuged immediately prior to ZnCl_2_ addition (t0) and after several time points, as specified in figure 4C. Pellets were solubilized in the appropriate volume of 2× β-mercaptoethanol gel-loading buffer (an OD_600_ of 1 gives 0.175 mg/ml of protein) to obtain a final concentration of 2 mg/ml and loaded onto SDS PAGE, using 4-12% precast gels (Thermofisher Scientific). Transfers were performed with iBlot2 transfer stacks (Invitrogen) and nitrocellulose membranes were incubated with the anti-CzcR (30), anti-penta HIS (Invitrogen) and anti-HSP70 (26) antibodies. Blots were revealed by chemiluminescence using the Amersham Imager 680 System.

### EMSA

The ability of Zur to bind the 5’-UTR DNA region of *czcR*, was determined by electrophoretic mobility shift assays (EMSA). The *znuB* gene, the 5’-UTR of *czcR*, wt or containing the mutated *zur* box-es (#mut1, #mut2 or #mut 1+2), were amplified by PCR from either the *P. aeruginosa* wt genomic DNA or from the *in vitro* synthetized mutated DNA (GeneArt, Thermo Fisher Scientific, Fig 3A). Primers used for PCR amplification are indicated in supplementary table S2. Amplicons were then purified on agarose gel. Binding assays were performed as previously described (4). Results were analyzed by staining the gel with 0.1% ethidium bromide and revealing with UV light using a NuGenius instrument.

### DNAse I Footprinting assays

The *czcR* promoter was amplified by PCR from the wild type *P. aeruginosa* genomic DNA with 5’ Fluorescein (6FAM) primer as indicated in supplementary table S2. DNA, with the plus or minus strand labeled independently, were gel purified. 50 ng of DNA fragments were mixed with or without 1 μM Zur in 40 μL of EMSA binding buffer (4) supplemented with 5 μM zinc. The reaction was incubated for 30 min at room temperature, followed by partial DNAse I digestion as described previously (12). The sequence of the pME6001-*czcRS* plasmid, determined with the same FAM-labeled primers and the Thermo Sequenase Dye kit (Thermofisher Scientific) was used to align the peaks. All fragments were analyzed by capillary sequencing (Microsynth AG, Switzerland) using ILS600 as a size standard, then peaks were assessed using PeakScanner2 software (Thermofisher Scientific).

### RNA extraction

Overnight cultures of the *P. aeruginosa* wt or mutant strains were diluted to an OD600 of 0.1 in M-LB complemented with 30 μM TPEN and cultured for 2h30 before being induced with 2mM ZnCl_2_. For RT-qPCR, 0.5 ml of culture was mixed with 1ml of RNA protect (Qiagen) immediately prior to metal addition (t0) and after several time points as indicated in the figures. The *P. aeruginosa* wt culture intended for the 5’RACE experiment was collected after 5h of 2 mM ZnCl_2_ induction. Total RNA extractions were performed as mentioned in previous publications (30). Briefly, pellets were resuspended in 100 μL Tris-EDTA buffer supplemented with 5 mg/mL Lysozyme (Fluka) and incubated at room temperature for 10 min. The following steps were carried out with the RNeasy mini kit (Qiagen), according to the manufacturer’s directives. Residual genomic DNA was removed by treating with 10 units/sample of RNAse-free RQ1 DNAse (Promega) for 1h at 37°C followed by phenol-chloroform extraction and ethanol precipitation. Total RNA was then resuspended in RNAse-free water.

### RT-qPCR

Quantitative RT-PCR were performed as previously described (31). Briefly 500 ng of total RNA were reverse transcribed using random primers (Promega) with the ImProm-II reverse transcriptase (Promega) according to the manufacturer’s instructions. Quantitative PCR was performed on a tenfold dilution of resulting cDNA and using the SYBR Select Master Mix (Applied Biosystem), with the primers listed in table S2. Results were analyzed as formerly mentioned (32) and standardized with *oprF*.

### 5’RACE

The transcription starts of the *czcR* and *czcC* genes, were determined using the Rapid amplification of cDNA-5’ends (5’RACE) kit (Roche). Briefly, 5 μg of total RNA from the *P. aeruginosa* wt were reverse transcribed using the specific primers sp1R for *czcR* mRNA or sp1C for *czcC* mRNA. Resulting cDNAs were purified with the Wizard SV Gel and PCR Clean-Up System (Promega) and tailing with dATP and TdT. Tailed-cDNA strands were then amplified with the dT-Anchor primer (Roche) and the specific primer sp2R or sp2C (for *czcR* cDNA and *czcC* cDNA respectively). A twenty-fold dilution of the first PCR was used as template for a second PCR with primers Anchor (Roche) and sp3R for *czcR* cDNA or sp2C for *czcC* cDNA. PCR products were then purified by agarose gel electrophoresis and subcloned into a pCR2.1 vector with the TA Cloning kit (Life Technologies) according to the supplier’s instructions. Plasmids isolated from *E. coli* clones were analyzed by sequencing. The transcription starts of *czcR* or *czcC* indicated in figure 2A correspond to the first nucleotide following the polyA tail.

### Experimental relevance and Statistical Data

Regarding the graph representations, the mean values of at least three independent experiments are indicated in the figures, along with the corresponding standard deviations. When specified, statistical analyses were performed according to the Student’s *t*-test and significance *p*-values were set at *p* ≤0.05(*) or *p* <0.001 (***). For other figures, data points represent the mean of triplicate values.

## Acknowledgments

This work was supported by the Swiss National Science Foundation grant 31003A_179336 for K.P., Eccellenza grant PCEFP3_203343 for M.V and Ambizione grant PZ00P3_180142 for D.G.

## Supplementary information

**Additional file 1: Figure S1. A)** DNAse I footprinting of *czcR* promoter (template strand) in the absence or presence or of 1 μM Zur protein. DNA fragments were analyzed by capillary electrophoresis. The two Zur boxes are indicated by the numbers 1 and 2 in red. The sequencing reaction is visible below the figure (seq1) and the sequence with the two boxes indicated in red is shown below the figure. **B)** Transcription start site (+1) of *czcR* and *czcC* determined by 5’RACE. The CzcR box, according to (15) is indicated in the box and the −35, −10 sequences were determined *in silico* using BPROM program (33).

**Additional file 2: Figure S2.** Growth curves of the wt PAO1 strain, the Δ*zur* mutant and the *ΔzurΔznuB* double mutant without or in presence of 2 mM ZnCl_2_ (Zn). Standard deviations of the triplicates are indicated.

**Additional file 3: Table S1.** Strains and plasmids used in this study

**Additional file 4: Table S2.** Primers used in this study

