## Supplemental data for "A Zur-mediated transcriptional regulation of the zinc export system"

#### Figure S1

A)

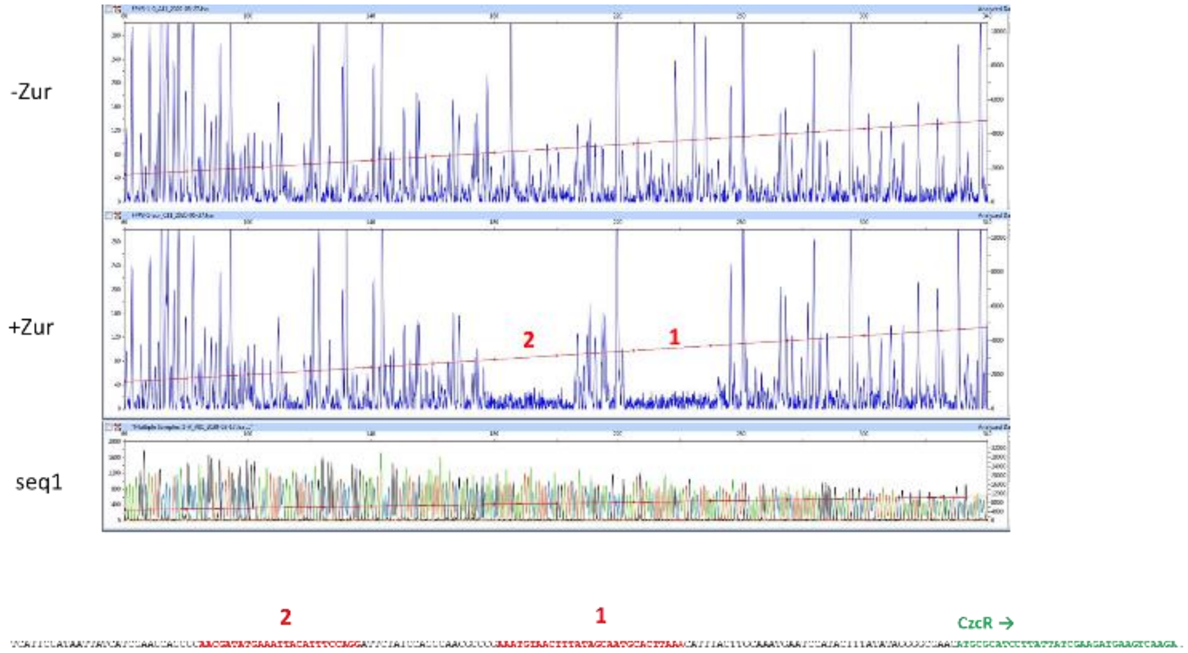

B)

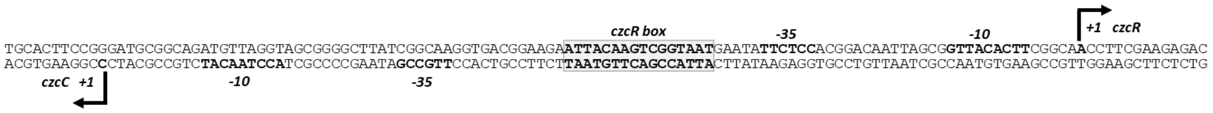

**Figure S1: A)** DNAse I footprinting of *czcR* promoter (template strand) in the absence or presence or of 1 µM Zur protein. DNA fragments were analyzed by capillary electrophoresis. The two Zur boxes are indicated by the numbers 1 and 2 in red. The sequencing reaction is visible below the figure (seq1) and the sequence with the two boxes indicated in red is shown below the figure. **B)** Transcription start site (+1) of *czcR* and *czcC* determined by 5'RACE. The CzcR box, according to (1) is indicated in the box and the -35, -10 sequences were determined *in silico* using BPROM program (2).

### Figure S2

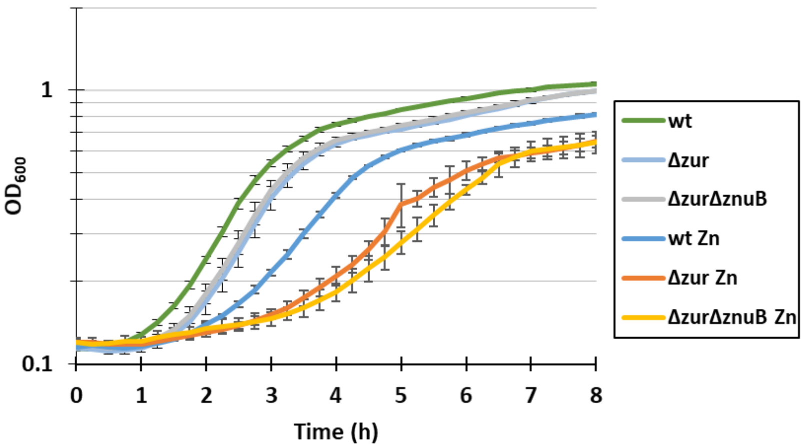

**Figure S2:** Growth curves of the wt PAO1 strain, the Δ*zur* mutant and the Δ*zur*Δ*znuB* double mutant without or in presence of 2 mM $ZnCl_2$ (Zn). Standard deviations of the triplicates are indicated.

### Table S1: Strains and plasmids used in this study

| | Strain or plasmid | Relevant characteristic(s)[a] | reference or source |
|---|---|---|---|
| ***P. aeruginosa*** | Wild type | PAO1 wild type | laboratory collection |
| | Δzur | PAO1 Δ*zur* | this study |
| | ΔczcRΔczcS | PAO1 Δ*czcR*Δ*czcS* | (3) |
| | ΔcadR | PAO1 Δ*copR*Δ*copS* | (4) |
| ***E. coli*** | DH5α | recA1, endA1, hsdR17, deoR, thi-1, supE44, gyrA96, relA1, Δ(lacZYA-argF), U169(ϕ80dlacZΔM15) | (5) |
| | BL21(DE3) | E. coli str. B F– ompT gal dcm lon hsdSB(rB–mB–) λ(DE3 [lacI lacUV5-T7p07 ind1 sam7 nin5]) [malB+]K-12(λS) | (6) |
| **Plasmids** | pME3087 | Suicide plasmid, Co1E1 replicon; $Tc^r$ | (7) |
| | pMMB66EH | Expressing vector carrying an IPTG-inducible promoter; $Ap^r$, $Cb^r$ | (8) |
| | pMMB66EH-*zur6HIS* | pMMB66EH derivative , carrying the *zur*gene fused with the C-terminal 6HIS-tag ; $Ap^r$, $Cb^r$ | this study |
| | pBBR1-*gfp* | Transcriptional *gfp* fusion cloning vector; $Ap^r$, $Cb^r$ | (9) |
| | pBBR1-*gfp* #1 | pBBR1 derivative , carrying the short *czcR* promoter; $Ap^r$, $Cb^r$ | this study |
| | pBBR1-*gfp* #2 | pBBR1 derivative , carrying the long *czcR* promoter; $Ap^r$, $Cb^r$ | this study |
| | pME6001-*czcRS* | pME6000 derivative , carrying the *czcR-czcS* operon under the *czcR* promoter; $Gm^r$ | this study |
| | pGex-2T-*zur* | GST-fusion expression plasmid,carrying the *zur* gene; $Ap^r$ | this study |

[a]Antibiotic resistance are indicated by r: Tc: tetracycline, Ap: ampicillin, Cb: carbenicillin, Gm: gentamicin.

#### Table S2: Primers used in this study

| | | | EMSA | | |
|---|---|---|---|---|---|
| Amplicon | PA number | Primer | Sequence 5'-3' | Position (from ATG) | Lenght |
| *pczcR* | PA2523 | 686 | GCCGGTACCACTTCGGCAACCTTCGAAGAG | -302 | 301 |
| | | 687 | GCCAGATCTGTTCGCCCCTATATAAAGTA | -1 | |
| *znuB* | PA5501 | 676 | CACCTCGCTGCTGATCATTC | 576 | 190 |
| | | 677 | AAAACTCAGCAGGAACAGGC | 765 | |
| | | | RT-qPCR | | |
| Amplicon | PA number | Primer | Sequence 5'-3' | Position (from ATG) | Lenght |
| *czcR* | PA2523 | 419 | GTCATCACCCGGACGCAGATCAT | 502 | 153 |
| | | 420 | GTAGCCGACGCCGCGAATGGTAT | 654 | |
| *czcC* | PA2522 | czcC1 | GGTCAGCATCGGCAGCAAGTACG | 834 | 206 |
| | | czcC2 | GGTCGTAGGCCTGTACCGCTTCG | 1039 | |
| *czcD* | PA0397 | 574 | GGCGTGGCCTTCTATATCCT | 285 | 183 |
| | | 575 | TCCAGACCTCCAGGTAGGC | 450 | |
| *cadA* | PA3690 | 1086 | CATCAACGCCCTGATGAGTA | 543 | 231 |
| | | 1087 | GCTTCCAGTTCCACTTGCTT | 773 | |
| *oprF* | PA1777 | 594 | GGTTACTTCCTGACCGACGA | 172 | 209 |
| | | 595 | TCGCTGTTGATGTTGGTGAT | 380 | |
| | | | DNA cloning | | |
| | Amplicon | Primer | Sequence 5'-3' | Lenght | |
| Footprinting | *pczcR* 5'FAM | 1271 | FAM-ACTTCGGCAACCTTCGAAGAG | 353 | |
| | | 857 | GGTGCAGGTAGTCGGCAGTC | | |
| | *pczcR* 3'FAM | 686 | ACTTCGGCAACCTTCGAAGAG | 353 | |
| | | 1272 | FAM-GGTGCAGGTAGTCGGCAGTC | | |
| GFP fusions | *pczcR*#1 | 686 | GCCggtaccACTTCGGCAACCTTCGAAGAG | 301 | |
| | | 687 | GCCagatctGTTCGCCCCTATATAAAGTA | | |
| | *pczcR*#2 | 895 | CGCggtaccGGAACCACGCAACCGTTCAT | 457 | |
| | | 687 | GCCagatctGTTCGCCCCTATATAAAGTA | | |
| Mutant | *Δzur* | 678 | CGgaattcCATGGTCGTGCTCGTCGTG | 510 | |
| | | 679 | caccagcgcgttgtccatcaggcgtccttctggtcGTGGTCATGGGGCTGGCA | | |
| | | 680 | GACCAGAAGGACGCCTGA | 500 | |
| | | 681 | CGggatccGTAGAGTTCAGCCTGGCC | | |
| *pMMB66EH-zur6HIS* | *zur* | 731 | GCCgaattcATGTACAAGATTGCGCCC | 500 | |
| | | 732 | GCCggatccTCAatgatgatgatgatgatgGGCGTCCTTCTGGTCCC | | |
| *pME6001-czcRS* | *czcRS* | 384 | GGGctcgagTCTGCTGATCGTCGTCGGCG | 2963 | |
| | | 385 | GGGaagcttGTTCTTCTCGCTGCCTGTTC | | |
| *pGEX2T-zur* | *zur* | 1044 | GCGggatccATGTACAAGATTGCGCCCAAGACCC | 503 | |
| | | 1045 | GCCgaattcTCAGGCGTCCTTCTGGTCCC | | |
| | | | 5'RACE | | |
| | | Primer | Sequence 5'-3' | | |
| | | sp1R | GCAGGCCGTCGATGCCATCG | | |
| | | sp1C | TGCCGCGCGCCGCTGGCGAT | | |
| | | sp2R | GGTGCAGGTAGTCGGCAGTC | | |
| | | sp2C | GCCGCCAGCTCGGGGTTGCT | | |
| | | sp3R | AGTCTTGACTTCATCTTCGA | | |
